# ADNP Modulates SINE B2-Derived CTCF-Binding Sites during Blastocyst Formation in Mouse

**DOI:** 10.1101/2023.11.24.567719

**Authors:** Wen Wang, Rui Gao, Dongxu Yang, Mingli Ma, Ruge Zang, Xiangxiu Wang, Chuan Chen, Jiayu Chen, Xiaochen Kou, Yanhong Zhao, Xuelian Liu, Hong Wang, Yawei Gao, Yong Zhang, Shaorong Gao

## Abstract

During early embryo development, the nuclear factor CTCF plays a vital role in organizing chromatin structure and regulating transcription. Recent studies have examined the establishment of nucleosome profiles around the CTCF motif sites shortly after fertilization. However, the kinetics of CTCF chromatin occupation in pre-implantation embryos have remained unclear. In this study, we utilized CUT&RUN technology to investigate CTCF occupancy in mouse pre-implantation development. Our findings revealed that CTCF begins binding to the genome prior to zygotic genome activation (ZGA), with a preference for CTCF anchored chromatin loops. Although the majority of CTCF occupancy is consistently maintained, we identified a specific set of binding sites enriched in the mouse-specific short-interspersed element (SINE) family B2, which are restricted to the cleavage stages. Notably, our data suggested that the neuroprotective protein ADNP may counteract the stable association of CTCF at SINE B2-derived CTCF-binding sites.

## Main

CCCTC-binding factor (CTCF) is a highly conserved, ubiquitously expressed DNA-binding protein with multivalent properties, including transcriptional regulation(Filippova et al. 1996; Ling et al. 2006), chromatin insulation(Bell et al. 1999; Cuddapah et al. 2009), genomic imprinting(Hark et al. 2000), and X chromosome inactivation(Chao et al. 2002). Recent evidence has also revealed CTCF’s role in organizing the three-dimensional (3D) chromatin architecture through the formation of short-range chromatin loops(Handoko et al. 2011; Dixon et al. 2012) or the novel phase separation behavior for long-range interactions(Lee et al. 2022; Wei et al. 2022). Composed of eleven zinc-finger (ZF) domains, CTCF recognizes DNA sequence diversity through the deployment of its distinct ZF clusters(Nakahashi et al. 2013). Sequence motif analysis has identified two major parts of the CTCF motif: the core, and upstream bound by ZF3-7 and ZF9-11, respectively. The core motif is present in most CTCF binding sites (CBSs)(Rhee and Pugh 2011), and the methylation status of the cytosine located inside the core motif can influence the binding affinity of CTCF(Bell and Felsenfeld 2000; Hashimoto et al. 2017).

Many studies have shown that CTCF plays a role during early mammalian development(Arzate-Mejia et al. 2018). Upon fertilization, terminally differentiated gametes are reprogrammed into totipotent cells through ZGA at the maternal-to-zygotic transition (MZT)(Lee et al. 2014; Schulz and Harrison 2019). After the first lineage segregation, pluripotent embryonic and extraembryonic trophectoderm lineages are generated(Hackett and Surani 2014). Appropriate CTCF occupancy in pre-implantation embryos is required for successful cell lineage segregation, as the absence of CTCF results in embryonic lethality in blastocysts(Wan et al. 2008; Chen et al. 2019; Andreu et al. 2022). In our previous study, we discovered nucleosome depletion regions (NDRs) around CTCF motif sites (CMSs) in both pronuclei, which suggested that CTCF may bind to the mouse genome prior to the timing of ZGA(Wang et al. 2022a). However, information about the dynamics of CTCF binding sites (CBSs) during early embryogenesis, particularly for pre-implantation embryos, is currently unavailable.

Transposable elements (TEs) are a rich source of diverse cis-regulatory regions for mammalian transcription regulation, including promoters, enhancers, and transcription factor binding sites(Sundaram et al. 2014; Trizzino et al. 2017; Hermant and Torres-Padilla 2021; Fueyo et al. 2022). In mouse, rodent-specific SINE B2 accounts for nearly one-third of CBSs(Bourque et al. 2008); while in humans, only 11% of CBSs are derived from TEs(Kunarso et al. 2010). Despite this difference, the 3D chromatin architectures are highly conserved between human and mouse(Harmston et al. 2017). Two models have been proposed to explain this discrepancy. The first model suggested that species-specific CMSs are suppressed by the activity-dependent neuroprotector homeobox (ADNP) protein and its cofactors to prevent unwanted CBSs(Kaaij et al. 2019a). The second model proposed that newly derived CBSs located near existing CBSs can act as a backup and cause no significant changes in 3D chromatin structures(Choudhary et al. 2020).

Here, we employed the recently developed Cleavage Under Targets and Release Using Nuclease (CUT&RUN)(Skene and Henikoff 2017) technique to produce CTCF binding profiles throughout early embryogenesis. We examined the intrinsic sequence features and epigenetic factors that impact CTCF re-binding events following fertilization. CTCF anchor sites (CASs) were preferentially established during embryogenesis. Furthermore, we identified over 1,000 cs-CBSs, which are highly enriched for SINE B2 repetitive elements, by comparing CBSs between 8-cell and blastocyst stage embryos. Our data suggested that ADNP may be responsible for the repression of these B2 SINE-derived cs-CBSs in blastocysts.

## Results

### CTCF re-binds to the genome prior to ZGA

To detect the dynamic of CBSs in early embryos, we utilized CUT&RUN technology to capture the genomic occupation of CTCF using low-input cells. We first generated CTCF CUT&RUN data in mouse embryonic stem cells (ESCs) using 1×10^5^ cells with two biological replicates (Supplemental table S1). The high reproducibility of CTCF CUT&RUN data was demonstrated by the highly correlated signal on potential CTCF binding sites (pCBSs) (Supplemental Fig. S1A, Supplemental tables S3 and S4, Pearson’s correlation coefficients = 0.94, pCBS definition see Materials and Methods for details). Next, we generated CTCF occupancy data using 5×10^3^ and 1×10^3^ cells with two highly reproducible biological replicates each (Supplemental Fig. S1A, Supplemental table S1, Pearson’s correlation coefficients ≥ 0.97). The signal of merged replicates from different input materials highly correlated on pCBSs (Supplemental Fig. S1B, Pearson’s correlation coefficients ≥ 0.89). We further identified CBSs through peak calling process on merged signal using MACS (version 2.1.3)(Zhang et al. 2008b; Feng et al. 2012). Most of the CBSs were identified in all three different input samples (Supplemental Fig. S1C, see Materials and Methods for details). Moreover, we performed *de novo* motif finding using MEME (version 5.0.5)(Bailey et al. 2006). The CTCF motif was ranked as the top motif for each sample (Supplemental Fig. S1D, see Materials and Methods for details). These results suggested that CUT&RUN can produce high-quality CTCF binding profiles using low input materials.

Subsequently, we generated CTCF occupancy data in mouse gametes and early embryos with two highly correlated biological replicates at each stage (Supplemental Fig. S1A and S2A, Supplemental table S1). In gametes, we identified thousands of CBSs that exist in the germinal vesicle (GV) but absent in the metaphase II (MII) oocytes (Fig. 1A, B). This phenomenon can be attributed to the differences in chromatin status across the cell cycle(Oomen et al. 2019). Following fertilization, we observed a resurgence of CTCF-binding signal at the pronuclei (PN)-3 stage (6 hours post fertilization, referred as 6 hpf), which is consistent with the well-organized NDRs around CMSs in both pronuclei observed in our previous work(Adenot et al. 1997; Wang et al. 2022a) (Fig. 1A, B). To confirm these CTCF re-binding events, we analyzed the dynamics of NDRs around PN-3 CBSs using our previous ultra-low-input MNase-seq (ULI-MNase-seq) data obtained from gametes and various pronuclei stage embryos(Wang et al. 2022a) (Supplemental table S2). Our analysis revealed more enhanced and evident NDRs around PN-3 CBSs compared to PN-3 non-binding CMSs in both pronuclei at 6 hpf, demonstrating that the reconstruction of CTCF binding after fertilization is reliable (Fig. 1C). NDRs around PN-3 CBSs were even detected in sperm rather than oocyte samples, and accordingly the nucleosome positioning was remodeled earlier in the male pronuclei after fertilization (Fig. 1C). This swift resurgence of CBSs following fertilization suggested that the binding of CTCF on chromatin is quite ubiquitous and fundamental in early embryos.

**Figure 1.**
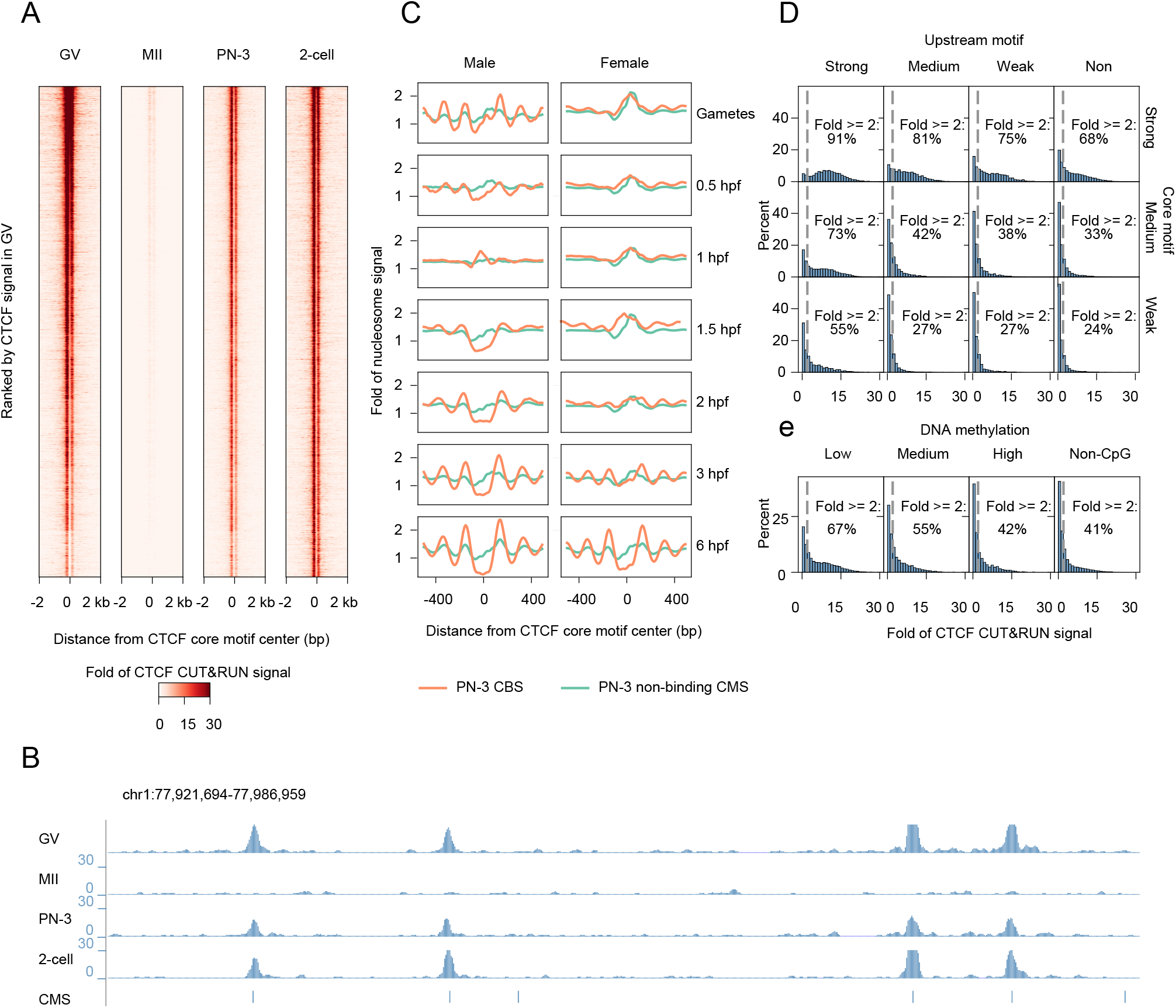
CTCF rebinds to genome upon fertilization. **A)** Heatmaps showing CTCF CUT&RUN signal in GV, MⅡ oocytes and PN-3, 2-cell embryos. CTCF binding were disappeared at MⅡ stage and largely recovered at PN-3 stage. GV: germinal vesicle, MII: metaphase II, PN: pronuclei. **B)** A snapshot of the browser view showing CTCF CUT&RUN signal within indicated region. CMS: CTCF motif site. **C)** Nucleosome profiles around CMSs at each indicated PN stage. Nucleosome depletion regions (NDRs) were established at male (left) and female (right) pronucleus as early as 1.5 hpf and 3 hpf, respectively. **D, E)** Histograms showing CTCF CUT&RUN signal in PN-3 embryos at CMSs grouped by motif strength (D) or DNA methylation status (E). The motif strength score of a CTCF-binding site was partitioned into strong (Top 25%), medium and weak (Bottom 25%) according to the motif occurrence within the site.

Previous studies have shown that CTCF binding can be influenced by multiple factors(Bell and Felsenfeld 2000; Rhee and Pugh 2011; Nakahashi et al. 2013). To determine the major factors involved in the CTCF re-binding process at the PN-3 stage, we first categorized the pCBSs into two groups, PN3-CBSs and PN3-non-binding CMSs, and measured various genetic features, including core and upstream motif scores, GC content, and CpG ratio, as well as epigenetic features such as DNA methylation, H3K4me3, H3K27me3, and H3K9me3 for each CBS group. We found the core motif score shows the highest predictive ability (AUROC= 0.73, Supplemental Fig. S2B). Besides, 91% of the pCBSs with both highest scores on core and upstream motifs (upper quarter) exhibited a CTCF binding signal (Fold of the CTCF CUT&RUN signal > 2) at the PN-3 stage (Fig. 1D). Conversely, the percentage reduced to 24% when the upstream motif was absent and DNA sequences had low core motif score (lower quarter) (Fig. 1D). Furthermore, lower DNA methylation levels in the core motif led to a higher percentage of CTCF binding signal present at PN-3 stage (Fig. 1E). The pCBSs lacking cytosine-phosphate-guanine (CpG) dinucleotides in their core motifs showed the lowest percentage of CTCF binding signal at PN-3 stage (Fig. 1E). Additionally, we observed a positive correlation between the PN-3 CBSs and H3K4me3, while a negative correlation was found between H3K9me3, H3K27me3 and the PN-3 CBSs (Supplemental Fig. S2C-E). These results demonstrated that reconstruction of CTCF binding is influenced by both intrinsic sequence and epigenetic features.

### CTCF anchor sites preferentially reconstruct after fertilization

After fertilization, 3D genome structures are reestablished, and the insulator function of these structures is primarily governed by CTCF(Guo et al. 2015; Du et al. 2017b). To investigate the maturation of these insulators, we computed the interaction probabilities of embryonic Hi-C data around the CTCF anchor sites (CASs) in ESCs (see Materials and Methods for details). The insulator function of CASs was found to increase at the 8-cell stage, as revealed by aggregated Hi-C signal analysis (Fig. 2A). Next, we investigated whether there were any discrepancies between CASs and non-CASs before the 8-cell stage. As expected, we observed that the rebinding of CTCF is established earlier on CASs than non-CASs (Fig. 2B, C), and the CASs exhibited a greater binding signal at each stage (Supplemental Fig. S3A). To elucidate the underlying reason driving the preference, we first assessed the CTCF motif scores for the above two anchor site groups, and found the core motif score on CASs is significantly higher than non-CASs (Fig. 2D).

**Figure 2.**
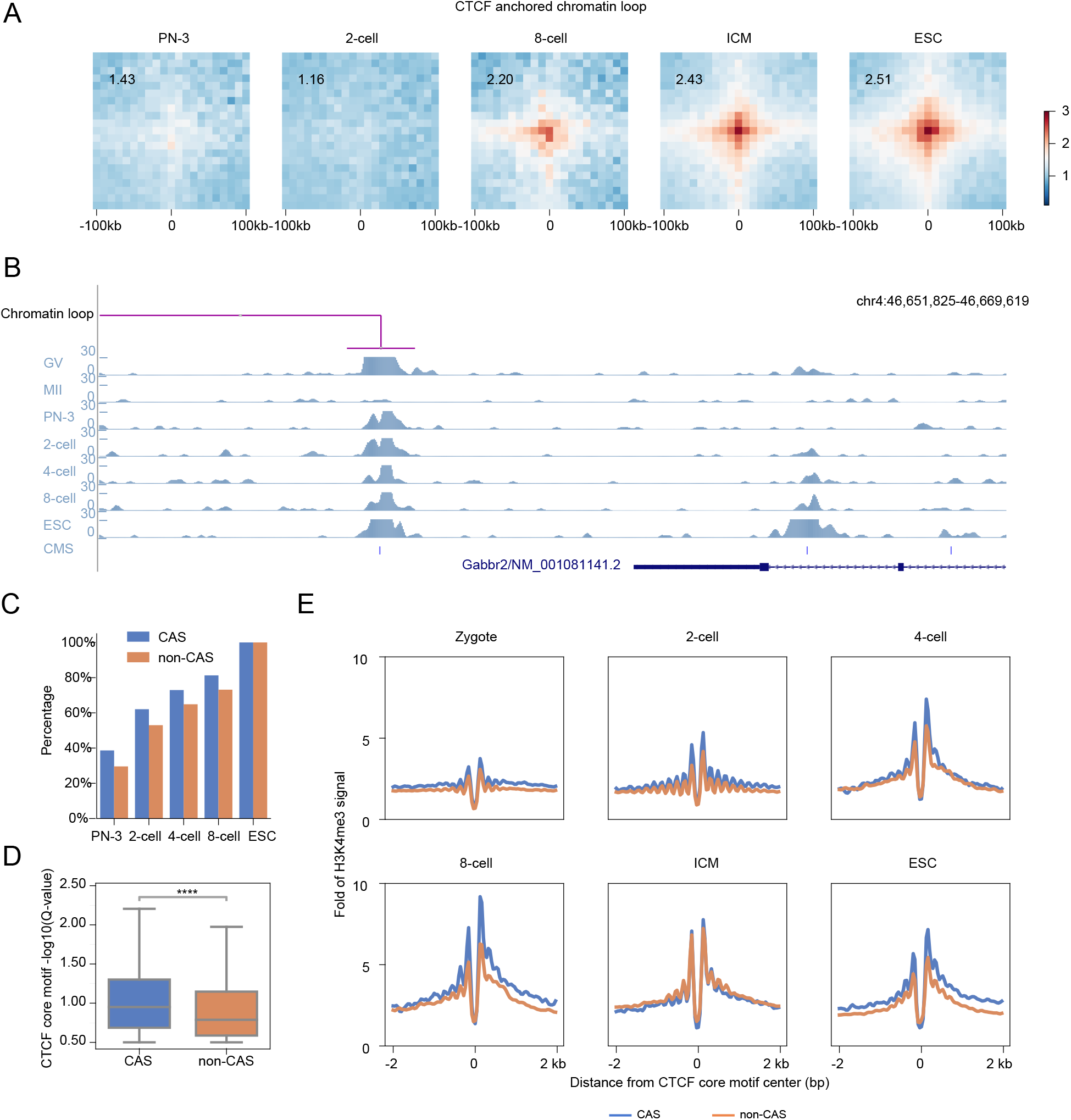
CTCF anchor sites preferentially reconstructs. **A)** Heatmaps showing the normalized average interaction frequencies at each embryonic stage for mESC CTCF anchor sites. The chromatin interaction was established from 8-cell stage. mESC: mouse embryonic stem cell, ICM: inner cell mass. **B)** A snapshot of the browser view showing CUT&RUN signal at indicated CTCF anchor sites and non-CTCF anchor sites. CAS: CTCF anchor site. **C)** Bar plots showing the binding percentage of distinct CAS classes in mESCs at each embryonic stage. CTCF anchor sites were established earlier than non-CTCF anchor sites. **D)** Boxplots showing that CTCF anchor sites exhibit stronger CTCF core motif than non-CTCF anchor sites. **E)** Line plots showing H3K4me3 signal levels around distinct CASs. CTCF anchor sites display higher H3K4me3 signals compared to non-CTCF anchor sites.

As insulators are characterized by multiple epigenetic features(Heidari et al. 2014), we next analyzed the dynamics of multiple epigenetic profiles at CASs and non-CASs, including DNase-seq, H3K4me3, H3K9me3, H3K27me3 signal and DNA methylation level. Our results showed that only H3K4me3 signal can clearly distinguish CASs and non-CASs before the 8-cell stage (Fig. 2E and Supplemental Fig. S3B-D). In summary, these findings suggested that core motif is closely related with the preferential reconstruction on CTCF anchor sites, and H3K4me3 may correlate with the reconstruction and maturation of CTCF-mediated insulators.

### Cleavage-specific CBSs are derived from SINE B2

Although most CTCF-binding sites remain stable during the 2-cell to blastocyst stage (Fig. 3A and Supplemental Fig. S4A), it has been revealed that loss of CTCF affects the efficiency of morula-blastocyst transition(Wan et al. 2008; Andreu et al. 2022). We then asked whether unique changes on CBSs occur during the first lineage segregation. We compared CBSs at the 8-cell embryos, inner cell mass (ICM) and trophectoderm (TE) at the blastocyst stage, and found thousands of CBSs lost in both ICM and TE cells (Fig. 3B, C). We defined this specific set of binding sites, which are restricted to the cleavage stages, as cleavage-specific CBSs (cs-CBSs); while the consistently maintained CBSs during the 8-cell to blastocyst stage are referred as reserved CBS (r-CBS) (Fig. 3B, C). To better validate the chromatin changes around these CBSs, we generated nucleosome occupancy data using ULI-MNase-seq(Wang et al. 2022a) at the corresponding stages to sample harvest for CTCF CUT&RUN-seq (Supplemental Fig. S4B-D, Supplemental table S1). Intriguingly, we found a 12 bp shift upstream in the NDR bottom around the cs-CBS, while the reserved CBS (r-CBS) had a subtler 4 bp shift upstream (Fig. 3D), suggesting that TF binding feature changes upon loss of CTCF binding on cs-CBSs. To verify if the occupation of CTCF on cs-CBSs uniquely happens in cleavage embryos, we collected 349 publicly available mouse CTCF ChIP-seq samples from Cistrome Data Browser(Mei et al. 2017) (Supplemental table S3, see Materials and Methods for details). CTCF binding events were rarely found in these cs-CBSs from all public mouse samples, indicating that the binding loss on cs-CBSs was permanent (Fig. 3E and Supplemental Fig. S4E). Next, we reviewed the binding kinetics of these two sets of CBSs. As shown, over half of r-CBSs obtained the CTCF binding in the PN-3 stage and this percentage was much higher than cs-CBSs until the 8-cell stages (Fig. 3F). Taken together, these results provided evidence for the existence of a specific set of cs-CBSs, which gains CTCF binding after ZGA but loses binding upon cell differentiation at the blastocyst stage.

**Figure 3.**
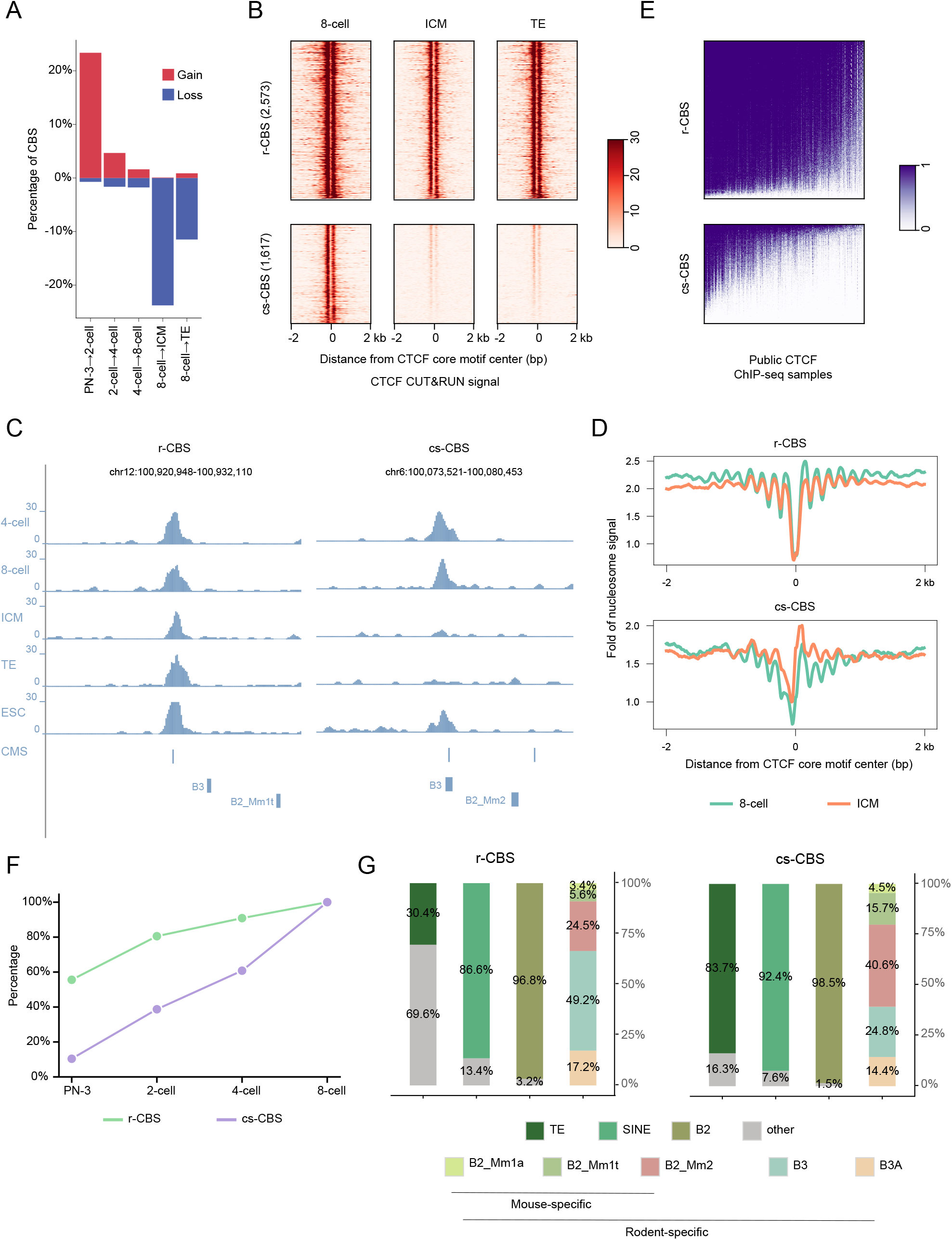
SINE B2 repeats introduce cleavage-specific CBS. **A)** Dynamics of CTCF binding sites (CBSs) during indicated embryo stages. **B)** Heatmaps showing CTCF CUT&RUN signal in 8-cell embryo and the ICM, TE lineages of blastocyst. ICM: inner cell mass, TE: trophectoderm. **C)** A snapshot of the browser view showing indicated r-CBS (left) and cs-CBS (right). r-CBS: reserved CBS, cs-CBS: cleavage-specific CBS. **D)** Nucleosome profiles around r-CBS (upper) and cs-CBS (lower) in 8-cell embryo and ICM. The bottom of NDR around cs-CBS shifted to upstream. **E)** Heatmaps showing the existence of CTCF signals in hundreds of publicly available mouse ChIP-seq samples. **F)** Line plots showing the step-wise establishment of distinct CBS classes. The majority of cs-CBSs were established during cleavage stages. **G)** Stacked barplots showing enrichment of each repeat family in r-CBSs (left) and cs-CBSs (right). TE: transposable element. TE annotation from UCSC Table browser (http://genome.ucsc.edu/cgi-bin/hgTables) was used.

We further investigated the influence factors, which correlated with the transient CTCF occupancy on cs-CBSs. Upon assessment of the phastCons scores of these CBSs, we observed that the cs-CBSs exhibited lower sequence convergence in comparison to the r-CBSs (Supplemental Fig. S4F). This temporary CTCF binding status echoes that in many species the CTCF motifs can be emerged from TEs, especially SINEs(Bourque et al. 2008; Schmidt et al. 2012), which was restricted by repressive histone modifications or competitive TFs(Kaaij et al. 2019a; Gualdrini et al. 2022). Therefore, we examined the enrichment of various kinds of TEs in cs-CBSs and r-CBS. Interestingly, the majority (76.2%) of cs-CBSs were derived from SINE B2, whereas only a minority (25.5%) of r-CBSs were derived from B2 (Fig. 3C, G and Supplemental Fig. S4G). Specifically, we found that cs-CBSs were enriched with SINE B2 when compared to r-CBSs, especially the mouse-specific B2 subfamilies (B2_Mm1a: 4.5% VS. 3.4%, B2_Mm1t: 15.7% VS. 5.6%, B2_Mm2: 40.6% VS. 24.5%, Fig. 3C, G and Supplemental Fig. S4G). These results suggested that cleavage-specific CBSs are mainly derived from SINE B2 elements.

### ADNP suppresses SINE B2-derived cleavage-specific CBSs

Previous work showed nearly 40% of CTCF binding sites in the mouse genome are derive from transposable elements(Sundaram et al. 2014), and CTCF binding in mammals is often associated with species-specific SINE B2 expansions(Schmidt et al. 2012; Rudan et al. 2015; Thybert et al. 2018). However, the detailed mechanisms enforcing the usage of these novel CTCF binding sites in early embryos is largely unknown. We identified a total of 61,454 SINE B2-derived CTCF motif sites and classed them in to three groups based on the distinct usage in pre-implantation embryos. Among them, 656 sites showed strong binding signals throughout the cleavage stage and inner cell mass/trophectoderm (ICM/TE) and were defined as B2-derived r-CBSs. Additionally, 1,232 sites exhibited loss of CTCF binding during blastocyst differentiation and were defined as B2-derived cs-CBSs. The majority of sites (21,260) showed no CTCF binding signals across embryonic stages and all publicly available samples, and were defined as B2-derived non-binding CMSs.

Subsequently, we explored potential factors contributing to the differential formation of these motif sites. We first assessed the sequence convergence among these three subgroups of B2-derived CMSs. B2-derived cs-CBSs showed a significantly lower degree of convergence compared to B2-derived r-CBSs, and B2-derived non-binding CMSs exhibited the lowest sequence convergence (Fig. 4A). Furthermore, the motif strength of these B2-derived CMSs exhibited a similar trend, with B2-derived non-binding CMSs displaying the weakest motif sequence and B2-derived r-CBSs displaying the strongest motif sequence (Fig. 4B and Supplemental Fig. S5A). Additionally, more than a quarter of B2-derived non-binding CMSs located in the inactive region-B compartments, while 14% of B2-derived cs-CBSs and less than 5% B2-derived r-CBSs were found in B compartments (Fig. 4C). These findings indicated that newly derived B2 SINE sequences with poor CTCF motif sequence were more likely to be retained but unused in the genome. Since sequences in A/B compartments contain different histone modifications(Rowley and Corces 2018), and H3K9 trimethylation in active chromatin was reported to restrict the usage of functional CTCF sites in SINE B2 elements, we reanalyzed publicly available histone modification data(Liu et al. 2016b; Wang et al. 2018b). The B2-derived non-binding CMSs generally exhibited moderate enrichment of any specific histone marks, while B2-derived r-CBSs displayed continuous enrichment of active H3K4me3 instead of repressive H3K9me3 modifications. Importantly, B2-derived cs-CBSs showed a transition from active H3K4me3 enrichment to repressive H3K9me3 enrichment, which coincided with the loss of CTCF binding across the blastocyst formation (Fig. 4D and Supplemental Fig. S5B). We also examined DNA methylation levels during the morula-to-blastocyst transition. However, the consistent demethylation observed in all three subgroups indicates that DNA methylation status is not responsible for the loss of CTCF binding in cs-CBSs (Supplemental Fig. S5C). These results indicated the loss of CTCF binding may correlate with histone modification rather than DNA methylation changes in B2-derived cs-CBSs during the first wave of embryonic cell differentiation.

**Figure 4.**
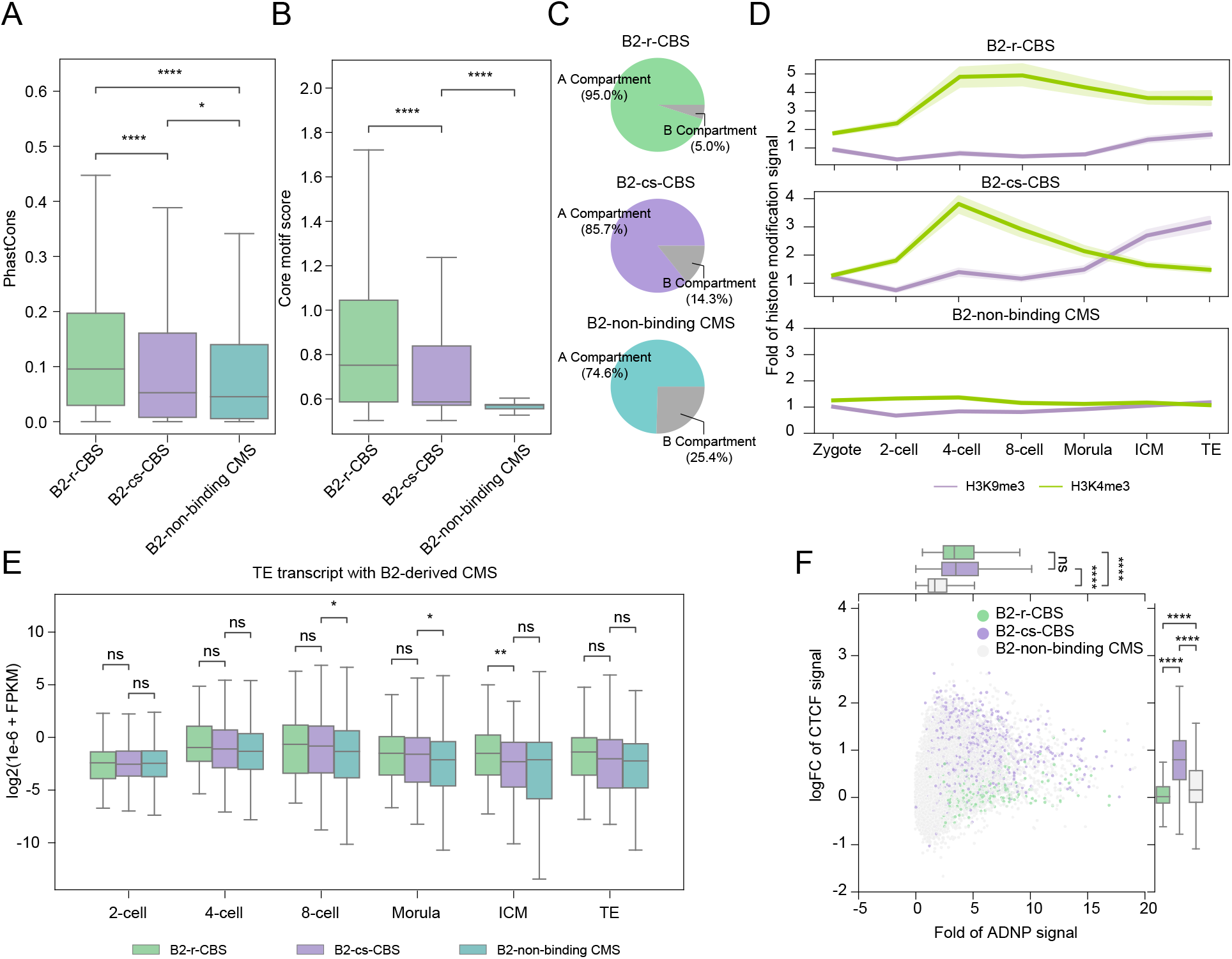
ADNP alters the CTCF binding at B2-derived cs-CSB in mESC. **A, B)** Boxplots showing the adjacent phastCons score (A) and core motif strength (B) of three distinct B2-derived CMS classes. B2-derived r-CBSs show the highest sequence convergence and strongest core motif, whereas B2-derived non-binding CMSs show lowest sequence convergence and weakest core motif. Motif strength was determined by −log10(q-value). **C)** Pie charts showing the percentage of distinct B2-derived CMSs in A/B compartment. **D)** Line plots showing active H3K4me3 and repressive H3K9me3 signals at three distinct B2-derived CMSs across the entire embryogenesis. B2-derived non-binding CMSs exhibit moderate enrichment of the indicated histone marks. B2-derived r-CBSs display sustained enrichment of active H3K4me3 modifications, whereas B2-derived cs-CBSs show a transition from active H3K4me3 to repressive H3K9me3 enrichment. **E)** Boxplots showing the expression levels of B2 repeats in distinct B2-derived CMSs at each embryonic stage. **F)** Joint plot showing the ADNP binding signal and log_2_-transformed CTCF ChIP-seq signal change after *Adnp* knockout in mESCs. Both B2-derived r-CBSs and B2-derived cs-CBSs show ADNP binding signal, however only B2-derived cs-CBSs show increased CTCF binding signal upon depletion of *Adnp*.

We also quantified the transcription levels of TEs contained within three different B2-derived CMS subgroups(Wang et al. 2018b) (see Materials and Methods for details), and observed a significant upregulation of SINE B2 in both B2-derived r-CBSs and B2-derived cs-CBSs compared to that in the B2-derived non-binding CMSs at both 8-cell and morula stages (Fig. 4E). However, as embryos differentiated into ICM and TE lineages, the expression level of SINE B2 in B2-derived cs-CBSs decreased to a level comparable to that of B2-derived non-binding CMSs, which was significantly lower than the expression of B2 in B2-derived r-CBSs (Fig. 4E). These findings demonstrated that loss of CTCF binding in B2-derived cs-CBSs during the embryonic cell differentiation is closely associated with establishment of H3K9me3 modifications and decreased transcriptional activity of corresponding TEs, but not correlated with DNA methylation status.

We then turned to investigate the suppression mechanism of B2-derived cs-CBSs in the blastocyst, and we noticed that the ChAHP complex was reported to counteract chromatin looping at SINE-derived CTCF sites in mouse cell lines(Kaaij et al. 2019a). In ChAHP complex ADNP is known to compete with CTCF especially at the motif sites in younger SINE elements. We reanalyzed publicly available mESC ADNP and CTCF ChIP-seq data(Kaaij et al. 2019a), and found the three subgroups of B2-derived CMS displayed different ADNP signal characteristics. Both B2-derived cs-CBSs and B2-derived r-CBSs showed more ADNP signal enrichment compared to B2-derived non-binding CMSs. However, only B2-derived cs-CBSs showed an increased CTCF signal upon *Adnp* depletion in mESCs (Fig. 4F and Supplemental Fig. S5B, D). Since ADNP can interact with HP1 and establish heterochromatin nanodomains (HNDs)(Ostapcuk et al. 2018a; Thorn et al. 2022), therefore we proposed ADNP may also be related with the increased H3K9me3 levels at B2-derived cs-CBS regions during the first lineage segregation (Fig. 4D and Supplemental Fig. S5B).

## Discussion

CTCF acts as a core 3D genome structure architecture protein. However, the gradual maturation of 3D genome structures raises the question about the present of CTCF binding in early embryogenesis(Du et al. 2017b; Ke et al. 2017). In this study, we generated the CTCF occupancy data from gametes to blastocysts and revealed the presence of CBSs in both pronuclei no later than PN-3 stage. Interestingly, this subset of PN-3 CBSs exhibits binding in sperms rather than oocytes, and is established earlier in the male pronucleus. This raises the intriguing question whether the timing discrepancy in CTCF acquisition between parental alleles is correlated to guidance provided by the original CTCF binding signals in both gametes, which warrants further investigation. Additionally, we observed although CTCF anchor sites can be established earlier, the formation of chromatin looping, as evident from current Hi-C data(Du et al. 2017b; Ke et al. 2017), appears to be significantly delayed. This suggests a multifunctional role of CTCF in early embryos, which may extend beyond the induction of 3D structure.

In mammals, transposable elements are major contributors of genetic material, ∼50% in mouse and human genome(Senft and Macfarlan 2021; Fueyo et al. 2022; Lawson et al. 2023). TEs participate in genome structure and transcription regulation through various mechanisms. Earlier studies in human and mouse revealed that a particular family of TEs is often over-represented in the set of binding sites for a given transcription factor. The TE-derived TF binding sites (TFBSs) and cis- or trans-chromatin regulation expand their regulatory potency and transcriptional activity to a wide range of cell types and developmental stages, which reflects an ongoing co-evolution that continues to impact mammalian development. Previous research has shown that SINE B2, a rodent-specific TE subfamily, introduces thousands of CTCF motifs into the mouse genome(Gualdrini et al. 2022). CTCF sites introduced by SINE B2 were found to play important role in chromatin structure in mESCs, and its binding activity is competed by the ChAHP complex (CHD4, ADNP, HP1), and inhibited by the enrichment of repressive histone modifications(Kaaij et al. 2019a). In early embryos, a large number of TEs, including SINEs, lose the repressive DNA or histone modifications and become reactivated, which provides an access for the widespread usage of TE-derived TFBSs. Therefore, investigating the generation and molecular dynamics of TE-derived TFBSs in early embryos is crucial for understanding the impact of TEs on development and evolution, and our study serves as an important example in this context.

Starting with the dynamic binding of CTCF on SINE B2-derived CMSs, we discovered distinct characteristics during embryogenesis (Fig. 5). Moreover, ADNP may be an important regulatory factor to govern the dynamic binding of CTCF to B2-derived cs-CBSs in early embryos, which is worthy of further experimental verification.

**Figure 5.**
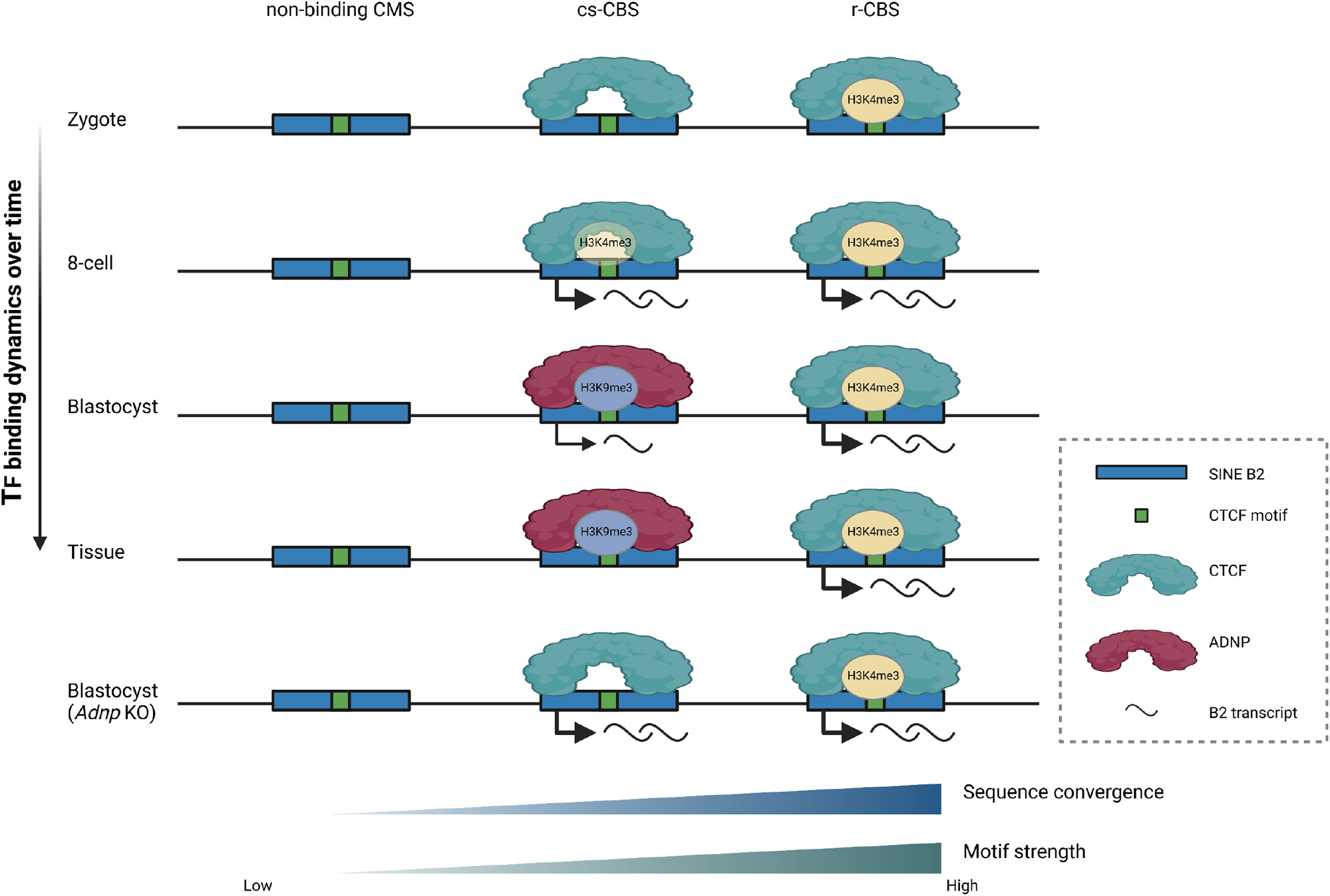
Schematics elucidate the fate of CMSs derived from SINE B2 expansions during embryogenesis (Created with BioRender.com). The youngest class in terms of evolution, inferred as B2-derived non-binding CMSs, exhibit neither CTCF binding nor histone marker enrichment. The B2-derived r-CBSs predominantly accumulate CTCF occupancy and H3K4me3, and exhibit high transcriptional activities, marked by highest sequence convergence and the strongest motif sequence. Alternatively, ADNP substitutes CTCF in the B2-derived cs-CBSs, resulting in transition from H3K4me3 to H3K9me3 and suppression of B2 expression during the morula-blastocyst transition.

Sequence motifs play a crucial role in determining TF binding sites (Fig. 1D and 2D). Misplaced or mis-activated TF binding sites, taking CBSs as an example, can disrupt the 3D structure of the genome and be associated with cancer(Katainen et al. 2015; Choudhary et al. 2020; Fang et al. 2020; Han et al. 2021; Choudhary et al. 2023). New motif sequences are generated through point mutations, small indels, or insertions. The expansion of TEs in the genome can provide new material for genetic evolution, which may also introduce mass of motif sequences to the embryonic genome(Gassler et al. 2022). However, introducing new CTCF motifs to the genome is also a challenging process, as it is crucial to avoid mistakes. Previous studies have primarily focused on the B2-derived CBSs and proposed that H3K9me3 and DNA methylation suppress these newly derived CBSs(Choudhary et al. 2020). When broadening our perspective to include all B2-derived CMSs, we observed that most of the B2-derived CMSs which lacking CTCF binding, termed as B2-derived non-binding CMSs, also lack of DNA methylation or H3K9me3 during early embryogenesis. Here, we proposed a model to elucidate the evolutionary process of B2-derived CMSs. The expansion of SINE B2 elements could introduce various types of new CMSs. B2-derived non-binding CMSs with a weak motif recognition represents a type of neutral mutation that remains compatible with the genome, whereas other types of B2-derived CMSs are mostly eliminated through nature selection. Subsequently, some of these B2-derived CMSs accumulate mutations and acquire improved motif recognition, sufficient for CTCF binding during the evolution. These sites may display different fates. Firstly, if it possesses the optimal CTCF motif, it becomes a r-CBS, serving as a backup for previous chromatin loop anchors or participating in the formation of new 3D genome structures. Secondly, if it contains a suboptimal motif, it becomes a cs-CBS. ADNP may counteract the stable localization of CTCF and suppresses these sites. Thirdly, if it possessed the optimal motif but forms harmful new 3D genome structures, it would get filtered through natural selection. Further discussion of this hypothesis will enhance our understanding of the evaluation of transcriptional regulatory networks.

## Materials and methods

### Animals and mouse oocyte or embryo collection

Specific-pathogen-free (SPF) mice were housed in the animal facility at Tongji University, Shanghai, China. All animal maintenance and experimental procedures were carried out according to the Health Guide for the Care and Use of Laboratory Animals and were approved by the Biological Research Ethics Committee of Tongji University.

To get pre-implantation embryos, 7-week-old C57BL/6N female mice were superovulated by intraperitoneally injection with 6 IU of pregnant mare’s serum gonadotropin (PMSG) and 6 IU of human chorionic gonadotrophin (hCG) (San-Sheng Pharmaceutical). The superovulated female mice were then mated with DBA2 male mice. Zygotes were collected from the oviducts of the female mice at 20 h after hCG injection, and were further cultured in G-1 PLUS medium (Vitrolife) to reach each corresponding developmental stage. PN3-stage zygotes were distinguished based on the microscopic observation of the size of the two pronuclei and the distance between them(Adenot et al. 1997). Then, we collected pre-implantation embryos at the following time points after hCG: 2-cell embryo, 36 h; 4-cell embryo, 58 h; 8-cell embryo, 71 h; morula, 85 h; blastocyst, 106 h. Germinal vesicle (GV) stage oocytes were collected 48 h after PMSG. To collect GV oocytes from 3-week-old mice, the whole ovaries were clipped mechanically with a razor blade. Fully grown GV oocytes were distinguished mainly based on the microscopic observation of the prominent but structurally homogenous bodies called “nucleolus-like bodies”(Inoue et al. 2007).

### Sample harvest for CUT&RUN-seq, total RNA-seq, uliNChIP-seq and ULI-MNase-seq

Samples of GV oocytes; PN3 stage, 2-cell stage and 8-cell stage; isolated inner cell mass (ICM) and trophectoderm (TE) of day 3.5 blastocysts or day 3.5 whole blastocysts; an ES cell line (R1) were harvested for CUT&RUN-seq, total RNA-seq and uliNChIP-seq. The zona pellucidae of the GV zygotes and cleavage-stage embryos were removed with 0.5% pronase E (Sigma) and the cleavage-stage embryos were then incubated in Ca^2+^-free Chatot-Ziomek-Bavister (CZB) medium for 10-20 minutes. Polar bodies were removed by gentle pipetting using a fire-polished glass needle. Single blastomeres were separated and manually picked before subjected to CUT&RUN library preparation and sequencing. For ICM and TE isolation(Liu et al. 2016b), the zona pellucidae of blastocysts were punched, followed by removal with 0.5% pronase E. The embryos were then incubated in Ca^2+^-free CZB for 20-30 minutes to disrupt the cell-cell junctions. ICM cells and TE cells were then distinguished and collected according to their sizes and shapes with the aid of a piezo-driven micromanipulator.

### Cell culture

The R1 ES cells were cultured on 0.1% gelatin pre-coated dishes in DMEM (Sigma) supplemented with 15% fetal bovine serum (FBS: Gibco), 1000U/mL leukaemia-inhibiting factor (LIF: Millipore), 1mM L-glutamine (Thermo Fisher Scientific), 100×non-essential amino acids (Millipore), 100×nucleosides (Sigma), 100×penicillin/streptomycin (Gibco), 0.1mM β-mercaptoethanol (Sigma), 1μM PD0325901 and 3μM CHIR9902. The ES cell line used in this study was regularly tested negative for mycoplasma contamination.

### CUT&RUN library construction and sequencing

CUT&RUN was conducted following the published protocols(Skene et al. 2018; Patty and Hainer 2021) with a few modifications. Briefly, all isolated fresh blastomeres or cells were washed three times with CUT&RUN wash buffer to avoid possible contamination and then transferred into a 1.5mL low-binding PCR tube (Eppendorf) containing 100µL of the wash buffer. The samples were then bound to Concanavalin A-coated magnetic beads (Bangs Laboratoris), which had been activated and resuspended in CUT&RUN binding buffer. After cell immobilization, bead-bound samples were successively incubated with appropriate amount of primary antibodies against protein-of-interest (CTCF antibody: Millipore, 07-729) in 50µL of CUT&RUN antibody buffer for overnight at 4°C. Cell membranes were permeabilized with 0.01% digitonin to allow the specific antibody to find its target. After unbound antibodies were washed away, 700 ng/mL protein A-MNase (pA-MN; a gift from Steven Henikoff lab) was added and incubated for 1 hour at 4°C. After washing, CaCl_2_ was added to a final concentration of 2 mM to activate pA-MN, and the digestion reaction was carried out for 30 min at 0°C and then stopped by adding 100μL 2× CUT&RUN stop buffer. The protein–DNA complex fragments were then released by 20-min incubation at 37°C. After transferring the supernatant to a new tube, 2µL of 10% SDS and 2.5µL of Proteinase K (Thermo) were added and incubated for 30 min at 55°C. DNA was then precipitated by phenol/chloroform/isoamylalcohol followed by ethanol precipitation with glycogen and then dissolved in nuclease-free water. Sequencing libraries were prepared using KAPA Hyper Prep Kit (KAPA biosystems) following the manufacturer’s instructions with slight modifications. Briefly, end repair was conducted for 30 min at 20°C followed by end-repair/A-tailing for 30 min at 50°C. After adaptor ligation for 30 min at 20°C, the DNA fragments were purified by 1.2×volume of AMPure beads (Beckman Coulter) followed by 18 cycles of PCR amplification with 2×KAPA HiFi HotStart Ready Mix. The final libraries were cleaned up with 1×volume of AMPure beads, and all CUT&RUN libraries were sequenced on a NovaSeq6000 (Illumina) platform at Novogene Co., Ltd. with paired-ended 150-bp reads.

### ULI-MNase library construction and sequencing

10-15 blastomeres or cells per replicate were isolated and washed before they were placed into 0.7 μL of lysis buffer (10 mM Tris-HCl, pH 8.5, 5 mM MgCl_2_, 0.6% NP-40) for individual reactions. Then, 2.5 μL of MNase master mix (MNase buffer, 0.125 U/μL MNase (NEB, M0247S), 2 mM DTT, and 5% PEG 6000) was added into each tube, and the reaction was incubated at 25 °C for 10 min for chromatin fragmentation. The reaction was stopped by the addition of 0.32 μL of 100 mM EDTA, and then 0.32 μL of 2% Triton X-100 was added to the reaction to release the fragmented chromatin. Then, 0.2 μL of 20 mg/mL protease was added, and the reaction was incubated at 50 °C for 90 min for protein digestion followed by incubation at 75 °C for 30 min for protease inactivation. The sequencing libraries were prepared using the KAPA Hyper Prep kit for the Illumina platform following the manufacturer’s instructions. After standard procedures including end repair and A-tailing, adapter ligation, post-ligation cleanup, and library amplification, the resulting products were subjected to a second round of PCR amplification with the same provided primers to generate sufficient DNA materials for high-throughput sequencing. Paired-end sequencing with 150-bp read length was performed on the HiSeq X Ten (Illumina) platform at Cloudhealth Medical Group Ltd.

### Generation of *Adnp* knockout mouse embryos

To ensure the effective deletion of Adnp in most embryos, two single guide RNAs (sgRNAs) targeting exon 4 and six sgRNAs targeting exon5 of Adnp gene were designed, respectively. The Cas9 mRNA and sgRNAs were produced as previously reported(Zhang et al. 2020). Sequence for each sgRNA was cloned into the sgRNA expression vector pUC57 and in vitro transcription was then performed using MEGAshortscript T7 transcription kit (Invitrogen). Meanwhile, Cas9 mRNA was *in vitro* transcribed using mMESSAGE mMACHINE T7 ultra transcription kit (Invitrogen). The integrity of manufactured mRNA was confirmed by electrophoresis. Both Cas9 mRNA and specific sgRNA were purified according to the standard protocol by phenol:chloroform extraction and ethanol precipitation, and then dissolved in nuclease-free water (Life Technologies). For microinjection, the Cas9 mRNA was diluted to 100ng/μL and each sgRNA mix was diluted to a final concentration of 50ng/μL. Zygotes were injected with approximately 10pL of Cas9 mRNA and sgRNAs using a Piezo-driven micromanipulator. Embryos were then observed and summarized from the 2-cell stage to the blastocyst stage. Besides, blastocyst stage embryos were harvested for CUT&RUN-seq, total RNA-seq and uliNChIP-seq. The genome targeting efficiency has been verified by PCR and Sanger sequencing. Primers for sgRNA are listed in Supplemental table S5.

### Reverse transcription and quantitative Real Time-PCR (RT-qPCR)

RNAs of 8-cell embryos were isolated using TRIzol Reagent and chloroform, with 1/10 volume of 3 mol/L NaAc and 1 μL glycogen was added to the aqueous phase of each sample. RNAs were then precipitated by isopropanol and washed twice with 75% ethanol before they were eluted with nuclease-free water. cDNA was then synthesized using All-In-One RT MasterMix (Applied Biological Materials). qPCR was carried out using TB Green Premix Ex Taq II (Takara Bio) and monitored by 7500 Fast Real-Time PCR System, and three technical replicates were performed for each sample. Relative expression level of target gene Adnp was normalized to the reference gene H2afz for embryo samples. qPCR primers for tested genes are also listed in Supplemental table S5.

### Total RNA library construction and sequencing

Embryos with zona pellucida and polar bodies removed were disrupted in TRIzol Reagent (Invitrogen) and total RNAs were isolated by chloroform extraction, coupled with isopropanol precipitation with 1/10 volume of 3M NaAc and 1 μL glycogen added to the aqueous phase of each sample. RNAs were washed twice with 75% ethanol before they were eluted with nuclease-free water. Purified RNAs were then subjected to library generation using SMARTer Stranded Total RNA-Seq Kit (Takara) following the manufacturer’s instructions. Briefly, random primers were used for reverse transcription, and the amplified cDNA was then subjected to ribosomal RNA depletion. Prepared RNA-seq libraries were sequenced on the Illumina NovaSeq 6000 platform with paired ends and 150-bp read lengths at Nanjing Jiangbei New Area Biophamaceutical Public Service Platform Co., Ltd.

### uliNChIP library construction and sequencing

uliNChIP-seq was performed as previously described(Brind’Amour et al. 2015) to capture the H3K4me3 and H3K9me3 status in embryos. Briefly, the harvested blastomeres or cells were subjected to nuclei extraction buffer. The MNase Master mix was then added and the mixture was incubated at 25°C for 10 minutes to allow MNase digestion. The reaction was stopped by 10 mM EDTA, and 0.1% Triton X-100 together with 0.1% DOC was added to lyse the nuclear membrane. The released chromatin was then diluted with ChIP buffer. After 1/20 volume of the reaction was saved as the input, the rest was incubated with primary antibody (H3K4me3 antibody: Cell signaling Technology, 9727; H3K9me3 antibody: Active Motif, 39161) -coated Dynabeads Protein A/G for overnight at 4°C. The ChIP samples were washed twice with low-salt wash buffer and twice with high-salt wash buffer. The washed beads were then incubated in hot elution buffer for 2 hours at 65°C with shaking. The eluted DNAs were further purified and subjected to sequencing library generation as described above. Paired-end 150-bp sequencing was also performed on ChIP libraries at Nanjing Jiangbei New Area Biophamaceutical Public Service Platform Co., Ltd.

### Immunofluorescent Staining

Embryos were fixed with 4% paraformaldehyde (Sigma) overnight at 4°C and then permeabilized with 0.5% Triton X-100 for 15 minutes at room temperature. The samples were blocked with 2.5% bovine serum albumin (BSA) (Sigma) at 25°C for 1 hour and then incubated with the primary antibodies against ADNP (R&D Systems), SOX2 (ABclonal) or CDX2 (Biogenex) overnight at 4°C. After washing three times with TBST, the samples were incubated with the appropriate secondary antibodies for 45 minutes. The nuclei were stained with 4’,6-diamidino-2-phenylindole (DAPI). All stained samples were observed using a Zeiss LSM880 confocal microscope. Images were processed and quantified in ImageJ software.

### Histone modification ChIP-seq data processing

Raw reads obtained were filtered using Trim Galore! (version 0.6.5, https://github.com/FelixKrueger/TrimGalore) with cutadapt (version 3.3)(Martin 2011) and the parameter “--trim-n”. The filtered reads were mapped to the mouse genome (mm10 assembly) using Bowtie 2(Langmead and Salzberg 2012) (version 2.4.2) with the parameters “--nomixed --no-discordant --no-unal”. Mapped reads with a mapping quality (MAPQ) score less than 30 were discarded and converted to BAM format using SAMtools (version 1.6)(Danecek et al. 2021). The BAM files were further converted to BED format using BEDTools (version 2.27.1)(Quinlan and Hall 2010). Replicates were merged, and the central 73 base pairs of all unique fragments were used for pile-up analysis. The resulting pile-up data was transformed into bigWig format for visualization and subsequent analysis using custom scripts and BEDTools (version 2.27.1)(Quinlan and Hall 2010). Broad H3K4me3 domains detection was performed as previously described.(Liu et al. 2016a)

### TF ChIP-seq data processing

ChIP-seq data for ADNP and CTCF were obtained from the Gene Expression Omnibus (GEO) datasets GSE97945(Ostapcuk et al. 2018b) and GSE125129(Kaaij et al. 2019b), respectively. Data were processed as previously described with mapping against mouse genome (mm10 assembly).(Wang et al. 2022b)

### WGBS data processing

Raw reads were obtained from the Gene Expression Omnibus (GEO) dataset GSE98151(Wang et al. 2018a) and processed as previously described with little modification(Liu et al. 2018). Raw reads were filtered using Trim Galore! (version 0.6.5, https://github.com/FelixKrueger/TrimGalore) with cutadapt (version 3.3)(Martin 2011) and the “--trim-n” parameter. The processed reads were then aligned to the mouse genome (mm10 assembly) using BSMAP(Xi and Li 2009). Methylation levels were determined using MCALL(Sun et al. 2014). Both BSMAP(Xi and Li 2009) and MCALL^9^ were from MOABS (version 1.3.9.6)(Sun et al. 2014). Subsequently, the methylation level of each CpG site and methylation signal tracks were generated for downstream analysis. Regions with mean DNA methylation levels below 0.2 were classified as low DNA methylation, those above 0.8 were classified as high DNA methylation, and the remaining sites were classified as medium DNA methylation.

### 3D chromatin-related data processing

We utilized publicly available CTCF ESC ChIA-PET loops from GSE99520(Weintraub et al. 2017) and performed a liftover to the mm10 genome assembly using CrossMap(Zhao et al. 2014). Raw reads of Hi-C data were obtained from GSE82185(Du et al. 2017a) and processed using HiC-Pro (version 3.0.0)(Servant et al. 2015) to extract all valid pairs with a minimum MAPQ greater than 30. The replicates were then merged and converted to the mcool format using cooler (version 0.8.2)(Abdennur and Mirny 2019). Aggregation peak analysis was conducted using coolpup.py (version 1.0.0)(Flyamer et al. 2020) on the previously mentioned CTCF loop anchors, with the following parameters: “--pad 100000” at a resolution of 10-kb. The mean of the center signal was assigned as the score.

### RNA-seq processing

To quantify gene expression, RNA-seq data were filtered using Trim Galore! (version 0.6.5, https://github.com/FelixKrueger/TrimGalore) with cutadapt (version 3.3)(Martin 2011), applying the “--trim-n” parameter. The filtered reads were then aligned to the mouse genome (mm10 assembly) using HISAT2 (version 2.1.0)(Kim et al. 2019) with the parameters “--no-mixed --no-discordant”. FPKM and read counts for each gene were calculated using StringTie (version 1.3.3b)(Pertea et al. 2015; Kovaka et al. 2019). To quantify the expression of TF transcripts, we followed the methods previously described(Shao and Wang 2021). The RNA-seq data were filtered using fastp (version 0.23.1)(Chen et al. 2018) with the parameter “-D” and then mapped to the mouse genome (mm10 assembly) using STAR (version 2.7.2a)(Dobin et al. 2013). Transcripts from various developmental stages (2-cell, 4-cell, 8-cell, Morula, ICM, and TE) were assembled from the mapping results using StringTie (version 2.2.1)(Pertea et al. 2015; Kovaka et al. 2019) and merged using Taco (version 0.7.3)(Niknafs et al. 2017). The expression of transcripts from all samples was determined by counting reads using featureCounts (version 2.0.1)(Liao et al. 2014), followed by redistribution using published scripts(Shao and Wang 2021) and normalization to FPKM. Transcripts overlapping with B2 were identified as B2 transcripts.

### ULI-MNase-seq data processing

The raw reads were subjected to quality control using Trim Galore! (version 0.6.5, https://github.com/FelixKrueger/TrimGalore) with cutadapt(Martin 2011) (version 3.3) and the parameter “--trim-n”. The filtered reads were then aligned to the mm10 genome using Bowtie 2 (version 2.4.2)(Langmead and Salzberg 2012) with the parameters “--no-mixed -- no-unal”. Reads with a mapping quality (MAPQ) score less than 30 were removed, and the resulting alignments were converted to BAM format using SAMtools (version 1.6)(Danecek et al. 2021). The BAM files were further converted to BED format using BEDTools (version 2.27.1)(Quinlan and Hall 2010). The replicates were then merged, and the central 73 base pairs of all unique fragments were used for pile-up. The resulting pile-up data was transformed into bigWig format for visualization and subsequent analysis, which was achieved using custom scripts and BEDTools (version 2.27.1)(Quinlan and Hall 2010).

### CTCF motif site (CMS) detection

The motif position weight matrix for CTCF (MA0139, MA1929, and MA1930) in MEME format was obtained from JASPAR(Castro-Mondragon et al. 2021). Motif sites were scanned against mouse genome (mm10 assembly) with MA0139.1 as previously described(Wang et al. 2022b). All CTCF motif sites were merged based on strand and location using BEDTools (version v2.29.2)(Quinlan and Hall 2010), and only the center positions were retained for further analysis. To determine the strength of the CTCF upstream motif, MA1929.1 and MA1930.1 were used to scan the 20bp-40bp region upstream of the CTCF core motif center against the genome sequence. CMSs overlapping with B2 by 10bp or more were classified as B2-derived CMS.

### CTCF CUT&RUN data processing

Raw sequenced read pairs were filtered by Trim Galore! (version 0.6.5, https://github.com/FelixKrueger/TrimGalore) with cutadapt (version 3.3)(Martin 2011) using the following parameters: “--trim-n”. The filtered reads were mapped back to the genome (mm10) using Bowtie 2 (version 2.4.2)(Langmead and Salzberg 2012) with the following parameters: “--no-mixed --no-discordant --no-unal”. Mapped read pairs (fragments) with MAPQ < 30 were discarded and converted to BAM format using SAMtools (version 1.6)(Danecek et al. 2021). BAM files were converted to BEDPE files using BEDTools (version 2.27.1)(Quinlan and Hall 2010). Replicates were merged, sampling down to 6M, and peak calling was performed by MACS (version 2.1.3)(Zhang et al. 2008a) with the following parameters: “-f BEDPE -g mm -q 0.05”. Peaks with fold ≥5 and Q-value ≤ 1×10−5 were maintained for the downstream analysis. The middle halves of fragments were piled up and transformed into bigWig format using custom scripts and BEDTools (version 2.27.1)(Quinlan and Hall 2010).

### CTCF binding site group definition

Publicly available CTCF ChIP-seq peaks were obtained from Cistrome Data Browser(Mei et al. 2016; Zheng et al. 2018) (Supplementary Table 4). These peaks were merged with embryonic CTCF CUT&RUN peaks and filtered based on motif overlap to identify potential CTCF binding sites (pCBS, Supplementary Table 5). Non-binding CMSs were CMSs that did not overlap with any pCBS. To determine the binding status of pCBS in each period, we assessed their overlap with peaks from the corresponding period samples (e.g., PN-3 CBS and PN-3 non-binding CMS, as shown in Fig 1). CTCF anchor sites (CAS) and non-CAS were CBSs in each period, depending on their overlapping situation with CTCF ESC ChIA-PET loop anchor(Du et al. 2017a).

The dynamics of pCBS were analyzed considering both the peak and signal. A pCBS was classified as a loss CBS if it overlapped with a peak and had a fold greater than 10 in the previous stage while was absent of peaks and had a fold less than 5 in the later stage. Conversely, a pCBS was considered a gain CBS if it overlapped with a peak and had a fold greater than 10 in the later stage while was absent of peaks and had a fold less than 5 in the previous stage. Specifically, the intersection of lost pCBS during 8-cell to ICM and 8-cell to TE were identified as cleavage-specific CBSs (cs-CBS), while CBSs with consistent binding in 8-cell, ICM, and TE with a signal greater than 10 were determined as r-CBS. Then, based on their overlap with B2, B2-r-CBS, B2-cs-CBS, and B2-non-binding CMS were defined.

### UCSC Genome Browser

The genome browser view was obtained using the UCSC Genome Browser(Kent et al. 2002) with Track data hubs(Raney et al. 2014) and visualized with smoothing with a mean of pixels. The signal was normalized by the average of the whole genome for visualization if not clarified.

### Statistical analysis

P-values were calculated by the two-sided Mann–Whitney U test if not clarified in figure legends. Asterisks represent statistical significance; (∗∗∗) P < 0.001, (∗∗) P < 0.01, (∗) P < 0.05, (n.s.) not significant.

### Data access

The raw sequence data reported in this paper have been deposited in the Genome Sequence Archive(Chen et al. 2021) in National Genomics Data Center(2023), China National Center for Bioinformation / Beijing Institute of Genomics, Chinese Academy of Sciences (GSA: CRA011730) that are publicly accessible at https://ngdc.cncb.ac.cn/gsa. A temporal link for reviewers is provided at https://ngdc.cncb.ac.cn/gsa/s/Z36BfD9k.

## Acknowledgments

We thank Dr. S. Henikoff for kindly sharing pA-MNase and advising CUT&RUN, and Hui Yang for suggestions on computational analysis. This work was primarily supported by the National Key R&D Program of China (2020YFA0509800 (R.G.), 2020YFA0113200 (Y.G.), 2022YFC2702200 (S.G.), 2021YFA1100300 (J.C.), 2019YFA0110100 (R.G.) and 2021YFA1302500 (Y.Z.)), the National Natural Science Foundation of China (31721003 (S.G.), 31922022 (Y.G.), 32270855 (R.G.), 32030022 (Y.Z.) and 31970642 (Y.Z.)) and the Natural Science Foundation of Shanghai Municipality (21ZR1465500 (R.G.)).

## Author contributions

R.G., W.W., Y.G., Y.Z. and S.G. conceived and designed the experiments. R.G., M.M., R.Z., and X.W. performed CUT&RUN experiments. R.G. and M.M. performed ULI-NChIP-seq and total RNA-seq experiments. W.W. and D.Y. performed the computational analysis. C.C, performed the ULI-MNase-seq. R.G., W.W., D.Y., M.M. and Y.G. designed and performed the data analysis. R.Z., J.C., X.K., Y.Z., X.L and H.W. assisted with the sample and library preparation. Y.H assisted with the manuscript preparation. W.W., R.G., D.Y, Y.G., Y.Z. and S.G. wrote the manuscript.

## Supplemental tables

Supplemental table S1. High-throughput sequencing data generated in this study.

Supplemental table S2. Public high-throughput sequencing data used in this study.

Supplemental table S3. Mouse public CTCF ChIP-seq dataset list for defining pCBS.

Supplemental table S4. Information of pCBS in this study.

Supplemental table S5. Primer sequences for sgRNA and RT-qPCR.

