## Supplemental Figures for "ADNP Modulates SINE B2-Derived CTCF-Binding Sites during Blastocyst Formation in Mouse"

Supplemental Figure S1

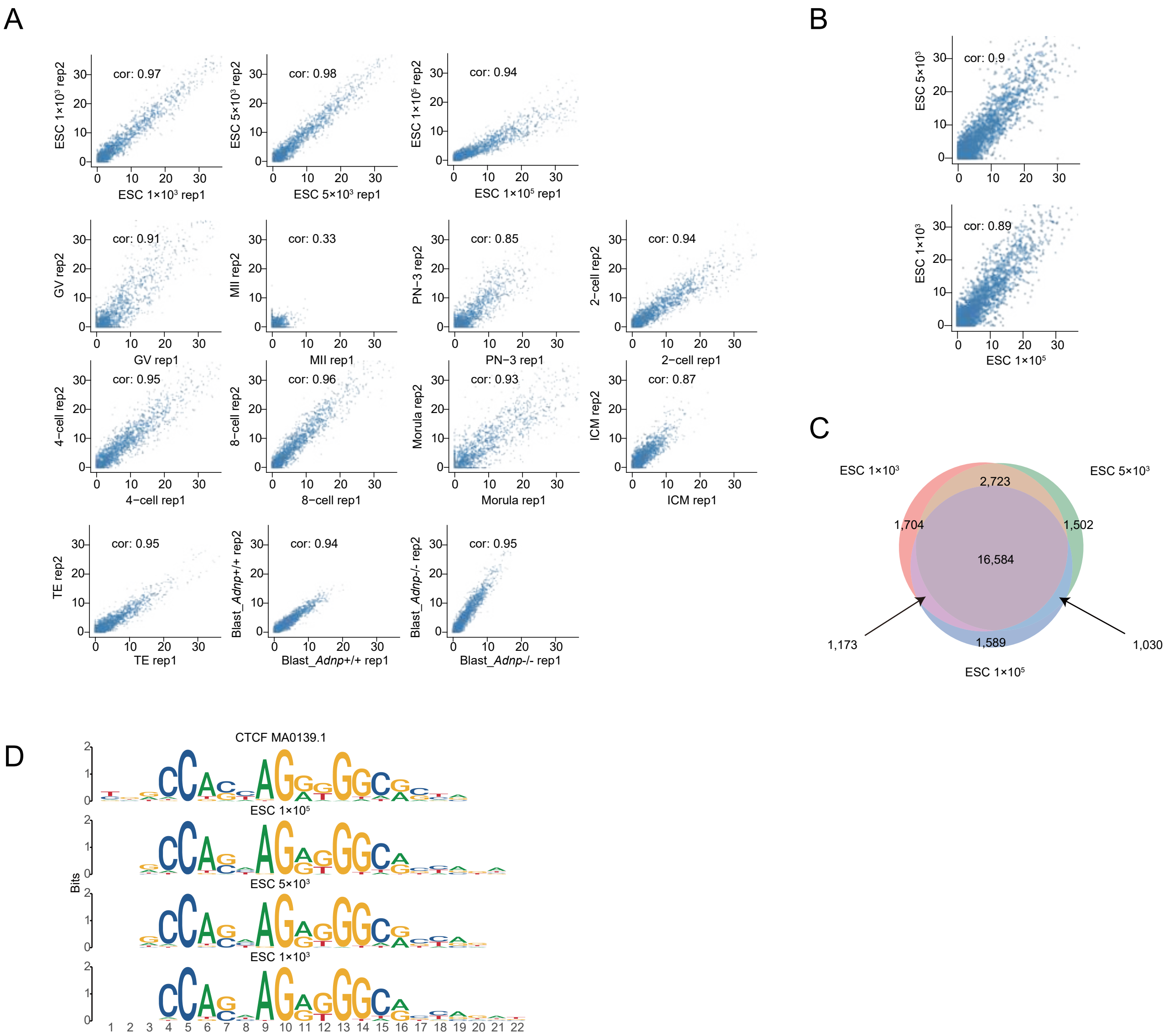

#### Supplemental Figure S1, Quality control of CTCF CUT&RUN data in mESCs and embryos.

**A)** Scatter plots showing high correlation of CTCF CUT&RUN signals on pCBSs between each replicate in mESCs and each embryonic stage. pCBS: potential CBS.

**B)** Scatter plots showing high reproducibility between CTCF CUT&RUN data on pCBSs generated from different input mESCs.

**C)** Venn diagram showing the peak overlap between CTCF CUT&RUN-seq data generated from different input mESCs.

**D)** *de novo* motif finding results based on CTCF CUT&RUN signals from different input mESCs.

### Supplemental Figure S2

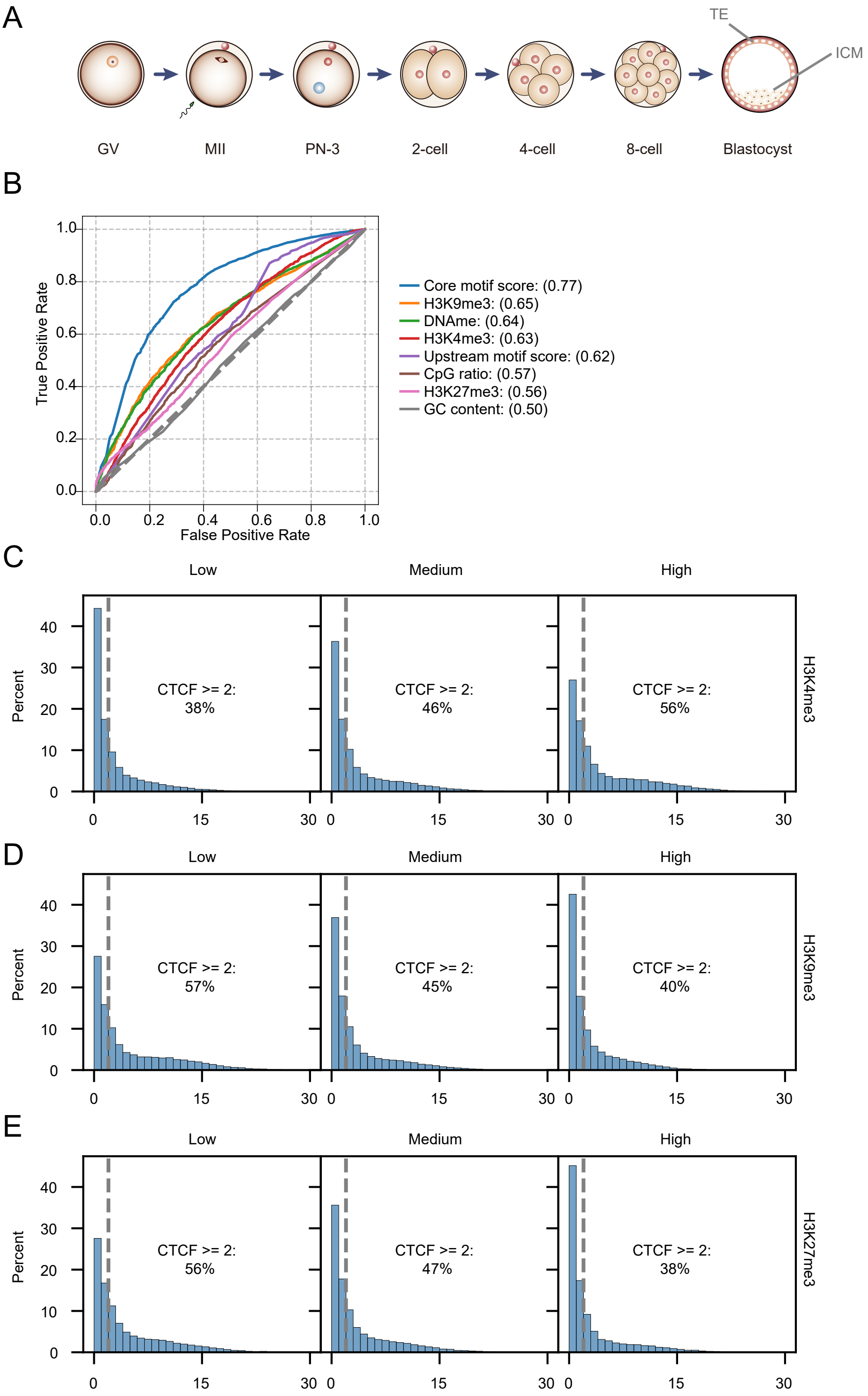

#### Supplemental Figure S2, Epigenetic features of CTCF binding within distinct CMSs at PN-3 stage.

**A)** Schematic for the gametes, pre-implantation embryos and cell line samples used in CUT&RUN analysis.

**B)** ROC curves showing the prediction power of each indicated feature in determining CTCF binding at PN-3 stage.

**C-E)** Histograms showing CTCF CUT&RUN signal in PN-3 embryos at CMSs grouped by H3K4me3 (C), H3K9me3 (D) and H3K27me3 (E) signals.

### Supplemental Figure S3

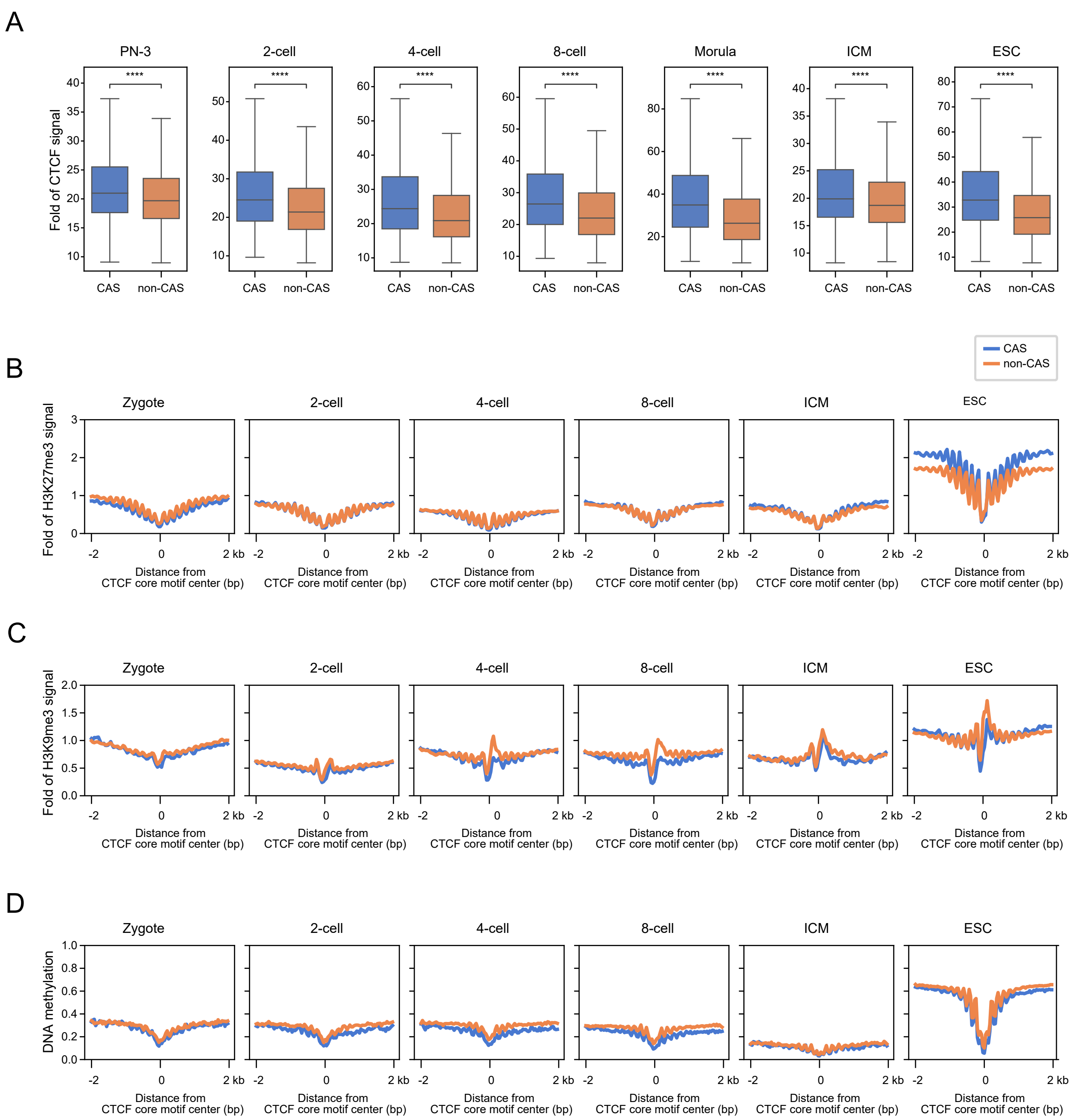

#### Supplemental Figure S3, Characteristics of CTCF enrichment and epigenetic states at CTCF anchor sites.

**A)** Boxplots showing CTCF CUT&RUN signal at CTCF anchor sites and non-CTCF anchor sites in each embryonic stage and mESCs. CTCF anchor sites exhibit stronger CTCF binding signals than non-CTCF anchor sites.

**B-D)** Line plots showing H3K27me3 (B), H3K9me3 (C) signals or DNA methylation status (D) around distinct CASs.

#### Supplemental Figure S4

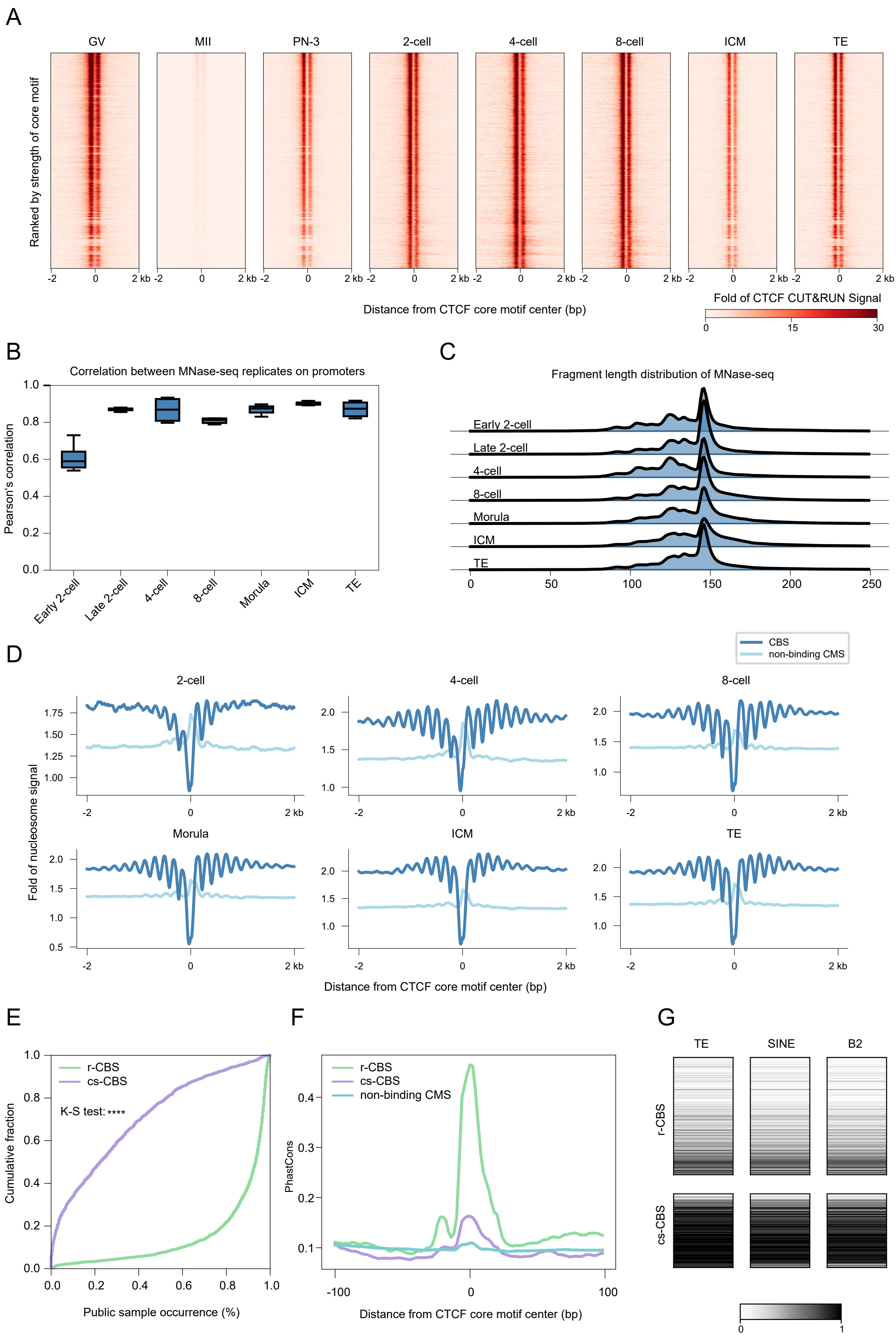

#### Supplemental Figure S4, cs-CBSs display low sequence convergence and high B2 enrichment.

**A)** Heatmaps showing the dynamics of CTCF CUT&RUN signal during mouse pre-implantation embryo development.

**B)** Boxplots showing the Pearson's correlation coefficients between MNase-seq replicates of each embryonic stage, which were calculated based on the nucleosome signal on promoter regions.

**C)** Density plot showing the length distribution of mapped fragments in MNase-seq libraries started from different embryonic stages.

**D)** Nucleosome profiles around CBSs and non-binding CMSs at each indicated embryonic stage.

**E)** Line plots showing the cumulative fraction of distinct CBSs corresponding to the binding ratios observed in publicly available CTCF ChIP-seq datasets.

**F)** Line plots deciphering the phastCons score around three classes of CBSs. cs-CBSs exhibited less sequence convergence in comparison to the r-CBSs. The profile of phastCons were download from UCSC (https://hgdownload.soe.ucsc.edu/goldenPath/mm10/phastCons60way/mm10.60way.phastCons.bw).

**G)** Heatmaps showing enrichment of indicated repetitive elements in r-CBSs (upper) and cs-CBSs (lower).

Supplemental Figure S5

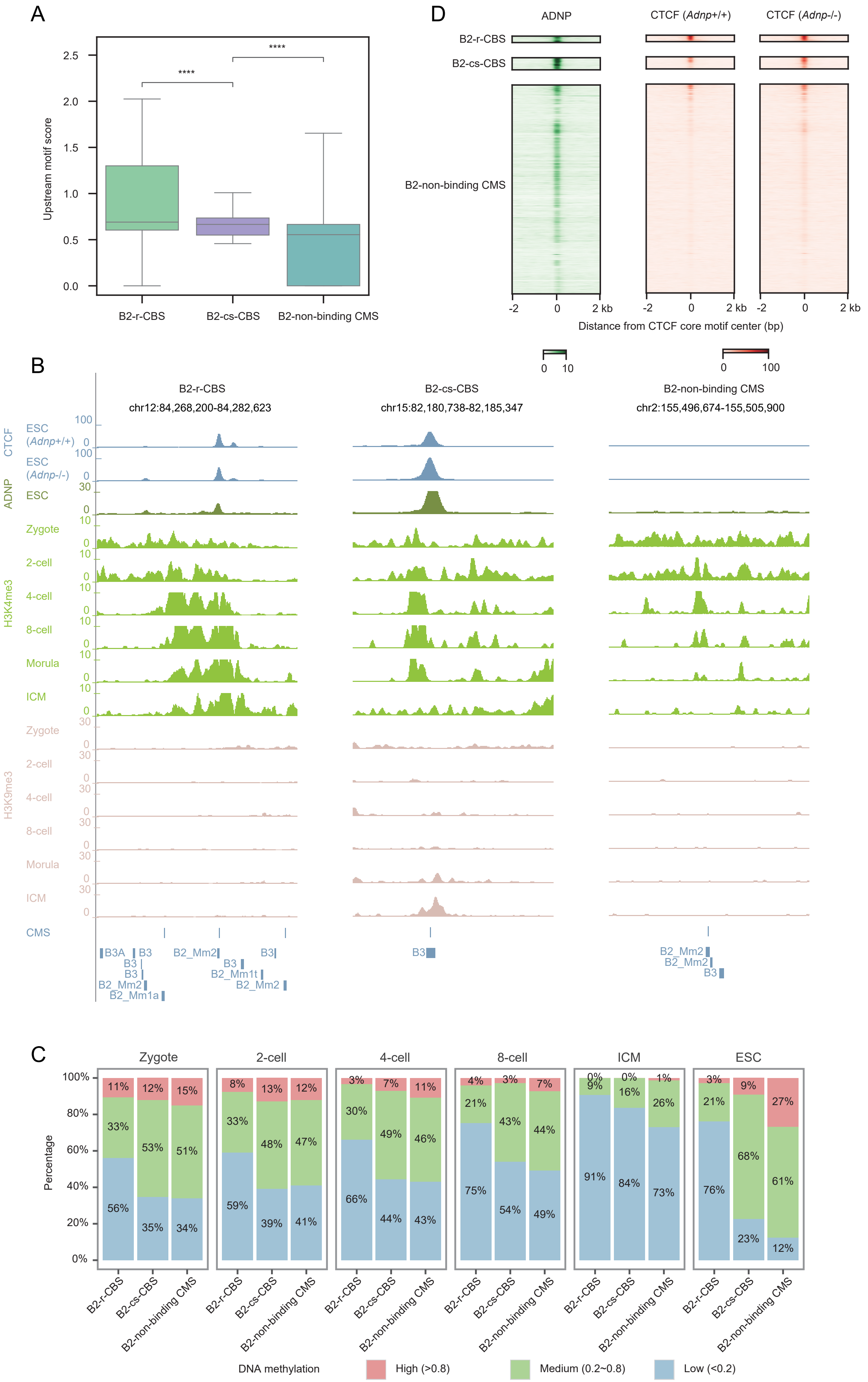

#### Supplemental Figure S5, CTCF signal and epigenetic states at distinct B2-derived CMSs upon depletion of *Adnp* in mESCs.

**A)** Boxplots showing the upstream motif strength around three classes of B2-derived CMSs. Motif strength was determined by -log10(q-value).

**B)** A snapshot of the browser view showing the CTCF, ADNP binding and histone modification changes at three representative distinct B2-derived CMSs in indicated cells and embryos.

**C)** Stacked barplots showing DNA methylation levels among distinct B2-derived CMSs in each embryonic stage and mESCs.

**D)** Heatmaps showing the ADNP binding signal and CTCF ChIP-seq signal changes at distinct B2-derived CMSs after *Adnp* knockout in mESCs.
